# Oncolytic Adenovirus Armed with cGAS Activates STING Pathway and Enhances Antitumor Immunity in Lung Cancer with Superior Combined Efficacy of PD-L1 Therapy

**DOI:** 10.64898/2026.01.15.699646

**Authors:** Qing-Wen Wang, Hua-Wei Xu, Yu-Sen Shi, Yi-Peng Zhang, Jie Jun, Dan-Ning Yue, Wei Zhao, Jia-Qiang Huang, Xiang-Lei Peng, Jie-Mei Yu, Jin-Sheng He, Yan-Peng Zheng, Yuan-Hui Fu

## Abstract

The extensive expression of STING in patients with non - small cell lung cancer (NSCLC) is closely associated with overall survival and other factors. Activation of the STING pathway can suppress NSCLC. However, the clinical translation of STING agonists remains hindered by challenges such as off-target effects, metabolic instability, and suboptimal pharmacokinetics. In this study, we engineered two oncolytic adenoviruses (OAds), OAd-HcGAS and OAd-McGAS, expressing human or murine cGAS, respectively, using an Ad5/3 chimeric adenovirus platform under regulation by the hTERT promoter to evaluatewhether OVs carrying the cGAS gene are capable of specifically activating the STING pathway within tumors and enhancing the anti - tumor efficacy of OVs both in *vitro* and *in vivo*.In vitro, OAd-HcGAS exhibited robust replication and potent cytolytic activity in tumor cells. It activated the STING–TBK1–IRF3 signaling axis, triggering a strong type I interferon (IFN-I) and pro-inflammatory cytokine response without compromising viral replication. In a murine Lewis lung carcinoma allograft model, intratumoral (i.t.) administration of OAd-McGAS led to substantial cGAS expression and consequential activation of the STING pathway. Moreover, the combination with anti-PD-L1 therapy resulted in tumor regression in over half of the cases. Notably, this armed oncolytic virus strategy enhanced the activation and infiltration of multiple immune cell populations. Collectively, these findings establish cGAS-expressing oncolytic adenoviruses as a novel and effective therapeutic strategy for lung cancer treatment.

**Graphical Abstract:** Viral replication & Transgene expression & Cancer treatment

## Introduction

Lung cancer remains one of the leading causes of cancer-related mortality worldwide^[1–4]^, with non–small cell lung cancer (NSCLC) accounting for approximately 85% of all cases^[2, 5–9]^. Surgical resection is the standard therapeutic option for early-stage NSCLC in eligible patients. However, patients diagnosed at late stages (IB to IIIB) hold a poor 5-year overall survival (OS) ranging from only 26% to 68%, with a subset progressing to metastatic disease^[10, 11]^. Beyond conventional approaches—including surgery, chemoradiotherapy, and targeted therapies—immunotherapy, particularly immune checkpoint inhibitors (ICIs), has become a mainstay of cancer treatment^[12–14]^. Despite the successful application of anti-PD-1, which has enabled long-term survival for some NSCLC patients^[15–17]^, approximately 70% of NSCLC patients still do not benefit from this treatment due to variations in PD-L1 expression levels^[18, 19]^.

Although ICIs have demonstrated significant efficacy across multiple solid tumors, they remain confined to a limited patient subset, largely due to the tumor microenvironment (TME)^[18, 20]^. In recent decades, driven by advances in genetic engineering and expanding applications of molecular biology, research on oncolytic viruses (OVs) has advanced rapidly, establishing them as a prominent strategy in lung cancer immunotherapy^[18, 21]^. Among various OVs, several, including oncolytic adenoviruses (OAds), are currently under investigation in clinical trials for a range of cancers^[22]^. OAds are characterized by high gene transduction efficiency, innate lytic capability, and ease of genetic modification^[23]^. Moreover, their cyclical replication and reinfection mechanism propagate the oncolytic effect to neighboring tumor cells, thereby amplifying therapeutic outcomes. A pivotal aspect of OAds therapy lies in their ability to induce immunogenic cell death, a process essential for priming antitumor immune responses^[24]^. Therefore, successful tumor elimination requires not only effective triggering of an immune response but also robust recruitment of immune cells.

As a major cytoplasmic DNA-sensitive mechanism, the cGAS-STING pathway has attracted increasing attention for its capacity to stimulate and link innate and adaptive immune responses, thereby contributing to antitumor defense. The cytosolic sensor cGAS recognizes double-stranded DNA (dsDNA) and, upon binding, undergoes a conformational change that enables the synthesis of the second messenger cGAMP^[25–28]^. Binding of cGAMP triggers STING oligomerization and recruitment of TBK1, which subsequently undergoes autophosphorylation and phosphorylates IRF3. Phosphorylated IRF3 then enables the nuclear translocation of IRF3 to drive type I interferon (IFN-I) expression, ultimately activating the innate immune system^[29, 30]^. Given the critical role of innate immunity in cancer immune surveillance, there is considerable interest in pharmacologically modulating innate immune pathways to enhance the efficacy of checkpoint inhibitors. The cGAS–STING pathway is frequently downregulated across multiple cancer types and is strongly associated with unfavorable clinical outcomes. Studies in lung cancer indicate that targeted activation of this axis can enhance antitumor immunity ^[31, 32]^. For instance, one study demonstrated that combining HSV-1-based OVs with 2’3’-cGAMP enhanced systemic antitumor immunity and immune memory responses in both local and distant tumors through dendritic cell (DC) activation, without compromising viral replication—even in tumor cells with impaired STING-IFN signaling^[33]^. Importantly, an intact STING pathway at the initial stage of tumorigenesis enables cGAS to detect self-DNA fragments leaking from the nucleus, thereby suppressing tumor progression^[34, 35]^. Although the STING pathway is frequently impaired in advanced tumor cells, this deficiency paradoxically creates a more favorable environment for OV replication^[33]^.

STING activation can be stimulated by diverse factors, such as ionizing radiation, DDRi, and STING agonists^[36]^. Although STING activation is critical for initiating innate immune responses, current STING agonists exhibit several limitations, including off-target activation, unstable metabolization, and suboptimal drug-like properties^[37]^. Moreover, given that STING is generally expressed in both tumor and normal cells, the constitutive pharmacological activation of STING agonists can cause adverse immune reactions^[38]^. These side effects significantly restrict their clinical application, highlighting the urgent need to develop strategies that achieve localized and tumor-specific activation of the STING pathway^[39]^.

To enhance the restoration of endogenous anti-tumor immune responses while directly activating antitumor immunity, we generated two 5/3 chimeric oncolytic adenoviruses (OAds) expressing species-specific cGAS and GM-CSF, designated OAd-HcGAS (human) and OAd-McGAS (murine). We first evaluated their ability to replicate, express transgenes, and elicit direct antitumor effects in lung tumor cells. Our in *vitro* results demonstrated that OAd-HcGAS effectively infected and lysed human NCI-H226 cells while concurrently activating the STING pathway. Furthermore, in a murine Lewis lung carcinoma (LLC) allograft model, i.t. administration of OAd-McGAS not only initiated the STING pathway and induced IFN-I production, but also enhanced lymphocyte infiltration and elicited a significant abscopal effect. This study indicates that cGAS-expressing oncolytic adenoviruses represent a promising approach for lung cancer therapy.

## Results

### Construction of Oncolytic Viruses Expressing Human and Murine cGAS

Analysis of the TIMER 3.0 database indicated broad expression of STING across both tumor and normal cells (Fig. S1A), highlighting the need for oncolytic viruses that selectively replicate in tumor cells to achieve tumor-restricted delivery of STING agonists. As illustrated in Fig. 1A, OAd-HcGAS and OAd-McGAS were constructed on a chimeric OAd-Z2 backbone, in which the E1B55k gene of the OAd-Z2 genome was fused with either human or murine cGAS via a P2A linker. The viruses were propagated through serial infection of HEK 293 cells, and transmission electron microscopy (TEM) confirmed typical adenovirus morphology (Fig. 1B). Compared with OAd-Z2 and the replication-deficient adenovirus H14 in HEK 293 cells, both OAd-HcGAS and OAd-McGAS exhibited comparable replication kinetics (*p* > 0.05) (Fig. 1C), indicating that the incorporation of cGAS and GM-CSF transgenes did not impair viral replication. Furthermore, ELISA verified GM-CSF expression (Fig. 1E), and Western blotting confirmed cGAS expression in NCI-H226 cells infected with OAd-HcGAS and in LLC cells infected with OAd-McGAS (Fig. 1F). These results collectively demonstrated the successful construction of novel recombinant oncolytic adenoviruses, OAd-HcGAS and OAd-McGAS, which express human and murine cGAS, respectively.

**Figure 1.**
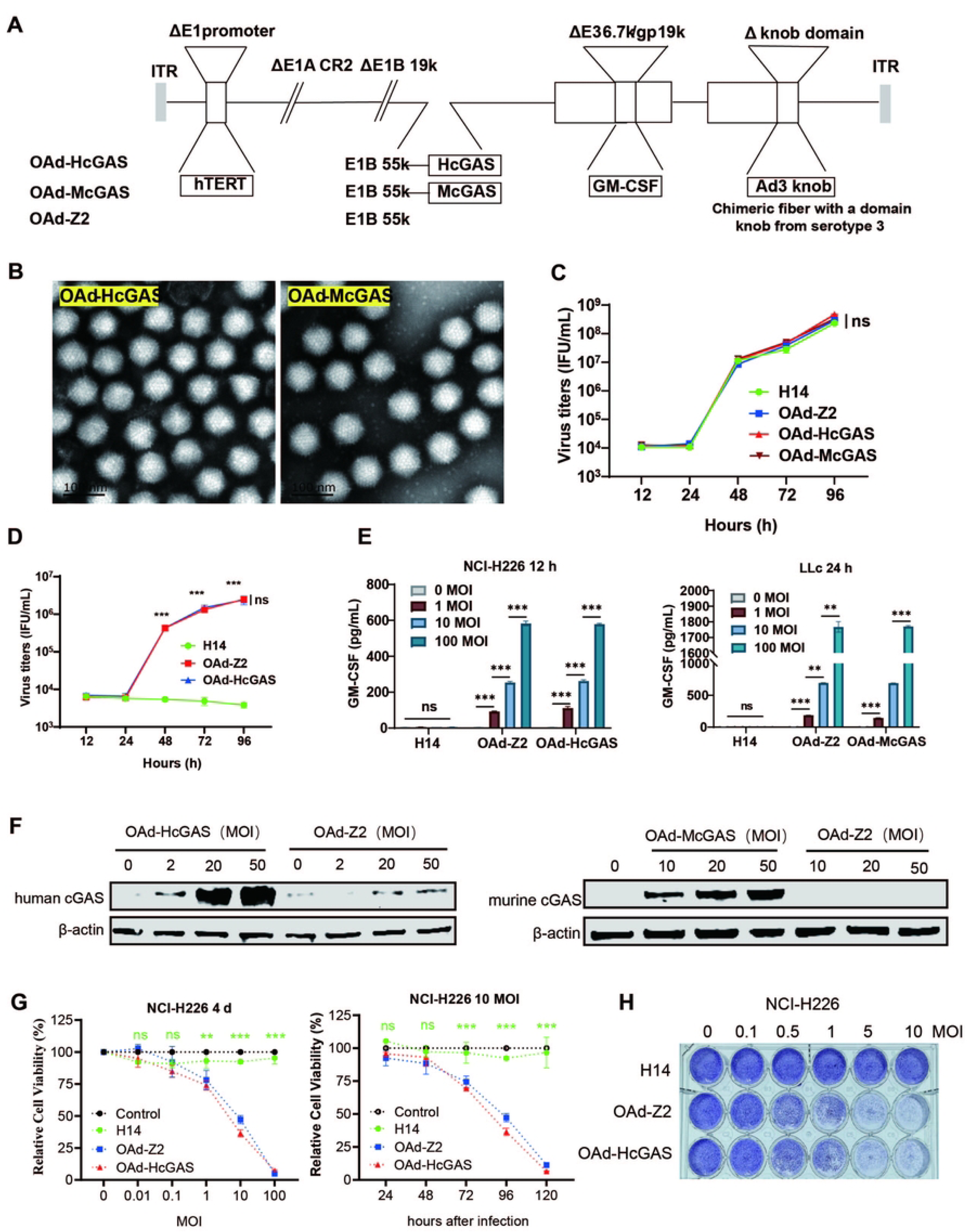
Functional verification of oncolytic adenovirus. (**A**) Schematic diagram of oncolytic adenovirus construction. OAd-HcGAS, recombinant adenovirus encoding human GM-CSF and human cGAS; OAd-McGAS, recombinant adenovirus encoding human GM-CSF and murine cGAS. **(B)** TEM of adenoviruses. **(C)** Viral replication capacity in HEK 293 cells infected at an MOI of 0.1 was measured using the plaque assay. **(D)** NCI-H226 cells were infected with recombinant adenovirus at an MOI of 0.5 and harvested at the indicated time points. Viral replication capacity determined by the plaque assay. **(E)** GM-CSF expression induced by OAd-HcGAS or OAd-McGAS was assessed in NCI-H226 and LLC cells. Cells were infected at various MOIs, and supernatants were collected 12 or 24 hours post-infection for GM-CSF quantification by ELISA. **(F)** cGAS protein expression in NCI-H226 and LLC cells after oncolytic adenovirus infection was evaluated via Western blotting. **(G)** The viability of NCI-H226 cells was evaluated using the CCK-8 assay at 24, 48, 72, 96, and 120-hours post-infection with OAds at varying MOIs, as well as at different time points following infection with a fixed dose of 10 MOIs. **(H)** Qualitative assessment of the cytocidal effect was conducted using the crystal violet assay at 96 hours post-infection across different MOI levels.Data are shown as mean ± SD. \*\**p* < 0.01, \*\*\**p* < 0.001, ns: no significant difference.

To assess the cytotoxic effects of OAd-HcGAS, various tumor cells were exposed to OAd-Z2 or OAd-HcGAS, and relative cell viability was measured using the CCK-8 assay. The results revealed a time-dependent decrease in cell viability across all tested cell lines at a multiplicity of infection (MOI) of 10. Moreover, viability further decreased in a dose-responsive manner 96 h post-infection as viral titers increased (Fig. 1G and S1B-1E). In parallel, crystal violet staining was performed to qualitatively assess tumor cell survival following infection at various MOIs. Both OAd-Z2 and OAd-HcGAS exhibited strong cytotoxic activity, nearly eliminating tumor cells at MOIs of 5 and 10 (Fig. 1H and S1B-1E). These findings indicate that OAd-HcGAS exhibits oncolytic efficacy comparable to or greater than OAd-Z2 across multiple tumor cell lines, demonstrating that the induction of antiviral immune responses does not impair its intrinsic lytic capability.

### OAd-HcGAS not only activates the cGAS-STING pathway but also does not compromise OAd-HcGAS replication in NCI-H226 tumor cells

Given that cytoplasmic dsDNA sensing by cGAS triggers STING signaling pathway^[40, 41]^, we next examined its capacity to induce STING pathway activation in NCI-H226 cells. As illustrated in Fig. 2A, infection of NCI-H226 cells with OAd-HcGAS at an MOI of 2 resulted in increased levels of phosphorylated TBK1 (pTBK1), which acts as the immediate downstream effector of STING activation. At elevated MOIs (20 and 50), despite a general reduction in total TBK1 and IRF3 expression, their phosphorylated forms (pTBK1 and pIRF3) accumulated substantially. These findings collectively demonstrate that OAd-HcGAS elicits a robust activation of the STING–TBK1–IRF3 signaling cascade in NCI-H226 cells, in contrast to the negative control OAd-Z2.

**Figure 2.**
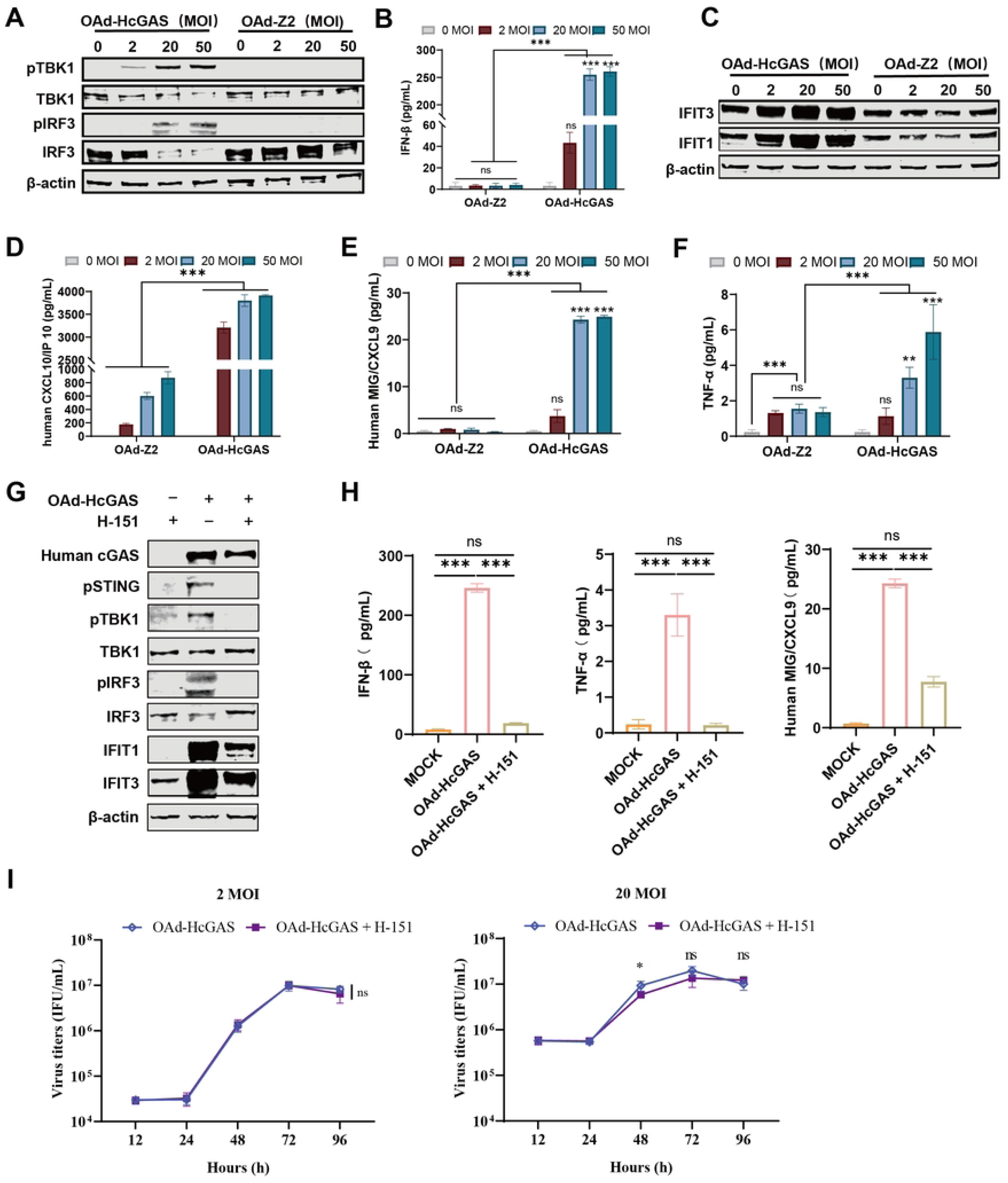
OAds activate the STING pathway in tumor cells. **(A)** Western blotting analysis of TBK1, pTBK1, IRF3, and pIRF3 in NCI-H226 cells infected with oncolytic virus at different MOIs for 48 hours. **(B)** ELISA quantification of IFN-β in cell supernatants. **(C)** Western blotting detection of IFIT1 and IFIT3 proteins. ELISA measurements of CXCL10 **(D)**, CXCL9 **(E)**, and TNF-α **(F)** in supernatants. **(G)** After STING pathway inhibition in NCI-H226 cells, infection with OAd-HcGAS (20 MOIs) was assessed by Western blotting for pSTING, TBK1, pTBK1, IRF3, pIRF3, IFIT1, and IFIT3. **(H)** ELISA analysis of IFN-β, CXCL9, and TNF-α secretion. **(I)** Viral replicative ability in NCI-H226 cells in the presence or absence of antagonist H-151. Data were shown as *X̄* ± *SD*, * *p* < 0.05, ** *p* < 0.01, *** *p* < 0.001, ns: no significant difference.

IFN-β is a principal downstream product of STING activation. To evaluate the capacity of OAd-HcGAS to initiate downstream signaling through this pathway, we examined the expression of IFN-β, proinflammatory cytokines (e.g., CXCL10), and interferon-stimulated genes, including IFIT1 and IFIT3, in NCI-H226 cells following OAd-HcGAS infection. As shown in Fig. 2B–2F and 2G–2H, OAd-HcGAS induced significantly higher levels of IFN-β, IFIT1, IFIT3, CXCL10, CXCL9, and CCL5 at different MOIs compared with OAd-Z2, which fails to activate cGAS–STING signaling in NCI-H226 cells. These findings suggest that OAd-HcGAS–mediated overexpression of cGAS effectively stimulates STING and subsequently activates the STING–TBK1–IFN signaling pathway in NCI-H226 tumor cells.

Efficient replication and proliferation of OVs in tumor cells, coupled with lysis of tumor cells by progeny virions, are central to their cytotoxic mechanisms. To assess the replication efficiency of OAd-HcGAS in NCI-H226 cells, viral titers were quantified at multiple time points following infection at an MOI of 0.1 using an immunoplaque assay. As shown in Fig. 1D, OAd-HcGAS replicated robustly in NCI-H226 cells, with no statistically significant difference compared to the control virus OAd-Z2. The STING pathway is widely recognized as a critical component of the cellular antiviral defense, and its activation is often associated with suppression of viral replication. To examine whether OAd-HcGAS–induced STING activation affects viral replication in NCI-H226 cells, cells were treated with 20 μM H-151 one hour prior to infection with OAd-HcGAS at MOIs of 2 and 20. Viral yields were then measured at 12, 24, 48, 72, and 96 h post-infection. As shown in Fig. 2I, STING pathway activation did not significantly suppress OAd-HcGAS replication in NCI-H226 cells.

### OAd-HcGAS activates and recruits immune cells *in vitro*

Infection with OAd-HcGAS markedly induced the secretion of CXCL10 and CXCL9 in NCI-H226 cells. To validate whether these secreted factors could promote lymphocyte migration, chemotaxis assays were performed using supernatants from OAd-HcGAS–infected NCI-H226 cells *in vitro*. As shown in Fig. 3A, supernatant derived from OAd-HcGAS–infected NCI-H226 cells significantly enhanced lymphocyte migration into the lower chamber compared to both the OAd-Z2 control and the blank group. These results demonstrate that OAd-HcGAS infection stimulates tumor cells to secrete chemokines that facilitate lymphocyte recruitment to the tumor site.

**Figure 3.**
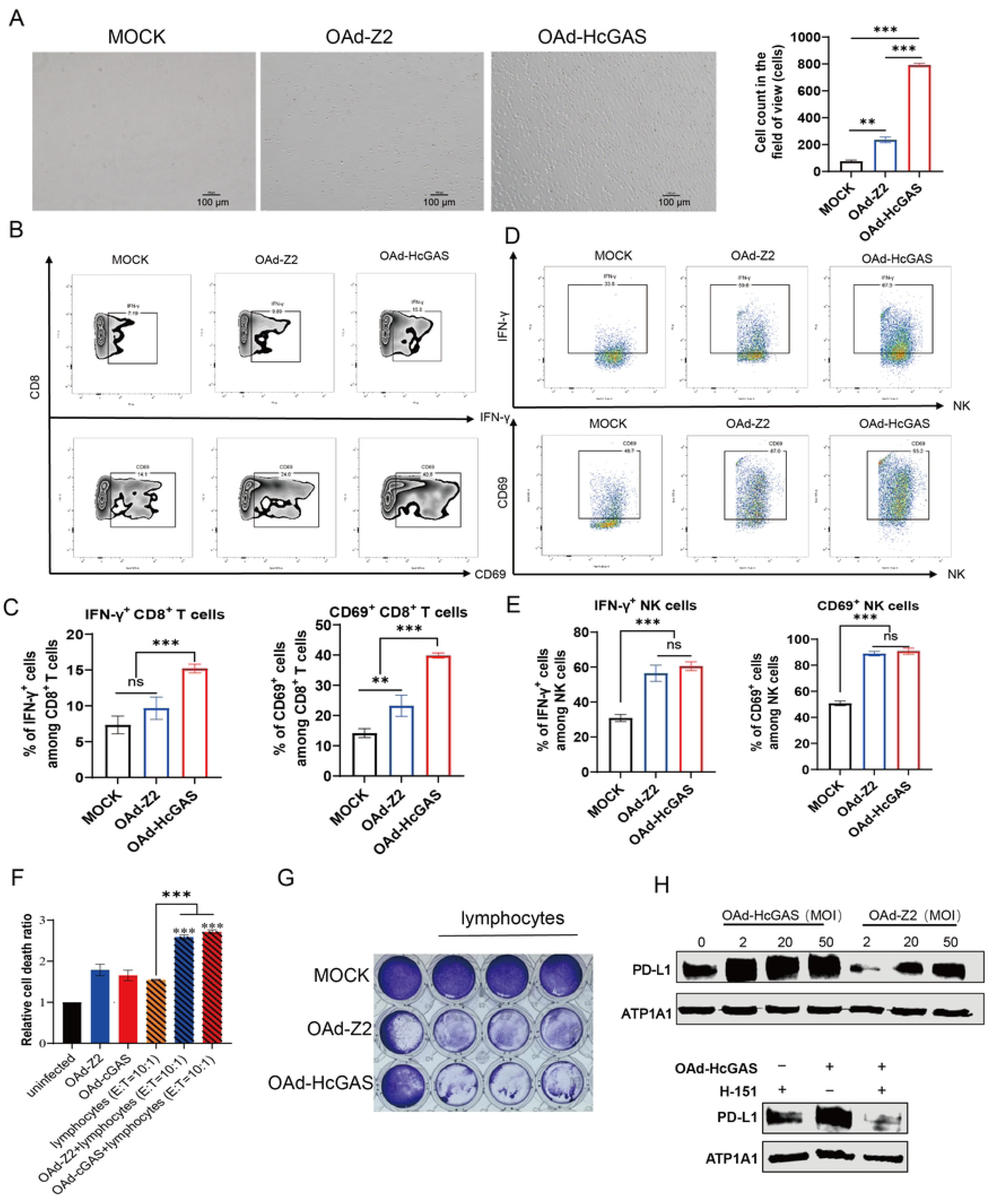
OAd-HcGAS can activate and recruit immune cells in vitro. **(A)** Lymphocyte recruitment and quantification in OAd-HcGAS-infected NCI-H226 cells. **(B-G)** NCI-H226 cells pre-infected with OAds were co-cultured with human PBMCs in a 1:10 ratio for 24 h before analysis by flow cytometry. Activation was measured by **(B, C)** expression of CD69 and IFN-γ on CD8 T cells or **(D, E)** expression of CD69 and IFN-γ on NK cells (N = 3 individual experiments). Cell death was measured by LDH cytotoxicity assay **(F)** and Crystal violet staining **(G)**. **(H)** PD-L1 proteins in NCI-H226 cells detected by Western blotting at 48 h after infection with OAd-HcGAS at different MOIs; Data were shown as *X̄* ± *SD*, ** *p* < 0.01, *** *p* < 0.001, ns: no significant difference.

To examine the immunostimulatory potential of OAd-HcGAS–infected NCI-H226 cells *in vitro*, a co-culture system was established combining these tumor cells with immune effector cells. As depicted in Fig. 3B–3E, co-culture with OAd-HcGAS–infected tumor cells led to a significant increase in CD69 expression on CD8^+^ T cells and NK cells (*p <* 0.05), a hallmark of early activation, and concurrently augmented IFN-γ secretion. Supernatants from the co-culture system were collected and subjected to an LDH assay to quantify LDH released into the extracellular space following tumor cell damage or lysis. As illustrated in Fig. 3F, the rate of tumor cell death was significantly higher in the co-culture system involving OAds (*p* < 0.05). Subsequently, the co-cultured tumor cells were subjected to crystal violet staining. As shown in Fig. 3G, NCI-H226 cells co-cultured with PBMCs and infected with OAd-HcGAS exhibited reduced staining compared to the control group. These findings suggest that OAds can effectively activate CD8^+^ T lymphocytes and NK cells to mediate cytotoxicity against tumor cells *in vitro*, with OAd-HcGAS exerting a stronger immune-stimulatory effect than OAd-Z2.

Additionally, consistent with previous reports^[42, 43]^, our study also demonstrated that OAd-HcGAS–infected NCI-H226 cells significantly upregulated PD-L1 protein expression (Fig. 3H), thereby providing a theoretical foundation for combining OAd-HcGAS with ICIs in subsequent therapeutic strategies.

### *In vivo* inhibitory effects of OAd-HcGAS on NCI-H226 lung cancer

To investigate the antitumor efficacy of OAd-HcGAS in vivo, a bilateral xenograft model was established by implanting NCI-H226 human lung cancer cells into BALB/c-nu immunodeficient mice. To evaluate the antitumor efficacy of OAd-HcGAS, the oncolytic adenovirus was directly injected into transplanted tumors of 100–150 mm³. By Day 12, OAd-HcGAS achieved superior tumor suppression (68.07 mm³, 65.81% inhibition) compared to OAd-Z2 (125.06 mm³, 39.02%) and PBS (215.9 mm³) in treated tumors. No significant difference was observed in untreated tumors among groups (Fig. 4B, 4C, 4G). Both OAd-Z2 and OAd-HcGAS resulted in significantly reduced tumor volumes on the treated side relative to the untreated side (*p* < 0.05), indicating localized therapeutic effects against NCI-H226 lung cancer.

**Figure 4.**
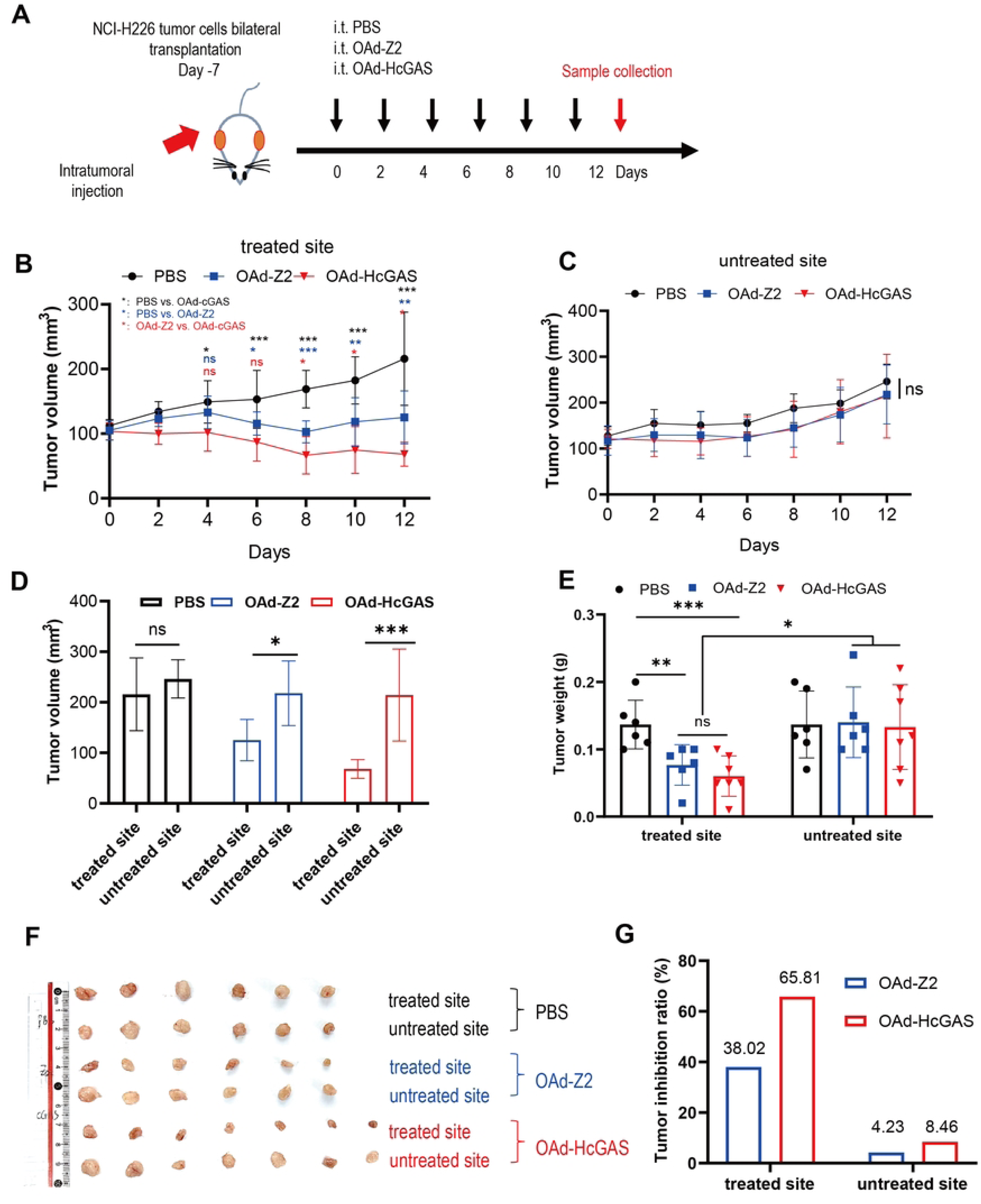
Inhibitory effect of OAd-HcGAS on NCI-H226 lung cancer. **(A)** Therapeutic scheme. In brief, one side was intratumorally administered with OAds (1×10^8^ PFU/tumor) every 2 days for 6 times in the bilateral subcutaneous NCI-H226 tumor model in BALB/c-nu immunodeficient mice, and the other side was untreated. **(B)** Tumor growth curves of treated tumors; **(C)** Tumor growth curves of untreated tumors; **(D)** Tumor volume, **(E)** Tumor weight, **(F)** Representative tumor images and **(G)** Tumor inhibition rate on Day 12 of the treated and untreated sites. Data were shown as *X̄* ± *SD*, * *p* < 0.05, ** *p* < 0.01, *** *p* < 0.001, ns: no significant difference.

### OAd-McGAS enhances antitumor efficacy and activates the cGAS-STING pathway *in vivo*

Bilateral LLC xenografts were established in C57BL/6N mice to evaluate tumor growth inhibition after unilateral i.t. injection. As shown in Fig. 5B, OAd-McGAS significantly reduced tumor volume on the treated side as early as Day 4 compared to OAd-Z2 (*p* < 0.05), whereas the reduction on the untreated side was modest and not statistically significant. By Day 10, tumor inhibition rates for OAd-McGAS reached 79.55% on the treated side and 58.24% on the untreated side, both exceeding those observed for OAd-Z2 (53.27% and 46.15%, respectively; Fig. 5C). Consistently, tumor weight measurements at Day 10 confirmed stronger suppression by OAd-McGAS, showing significant reductions compared to PBS on both sides (*p* < 0.05) and to OAd-Z2 on the treated side (*p* < 0.05; Fig. 5D). Notably, across multiple animal experiments, complete tumor regression was repeatedly observed at the treated site. Ki-67, a widely used marker of tumor growth and malignancy, was assessed by immunohistochemistry in Day 10 tumor tissues (Fig. 5G). Compared with the PBS and OAd-Z2 groups, Ki-67 expression was significantly decreased in the OAd-McGAS group on both treated and untreated sides (*p* < 0.05). These results indicate that OAd-McGAS, expressing murine cGAS, effectively inhibits the growth of LLC lung cancer xenografts. An even more encouraging finding is that complete tumor regression was observed at the treated site on day 8 in the OAd-McGAS treatment group during repeated animal experiments (Fig. S2F).

**Figure 5.**
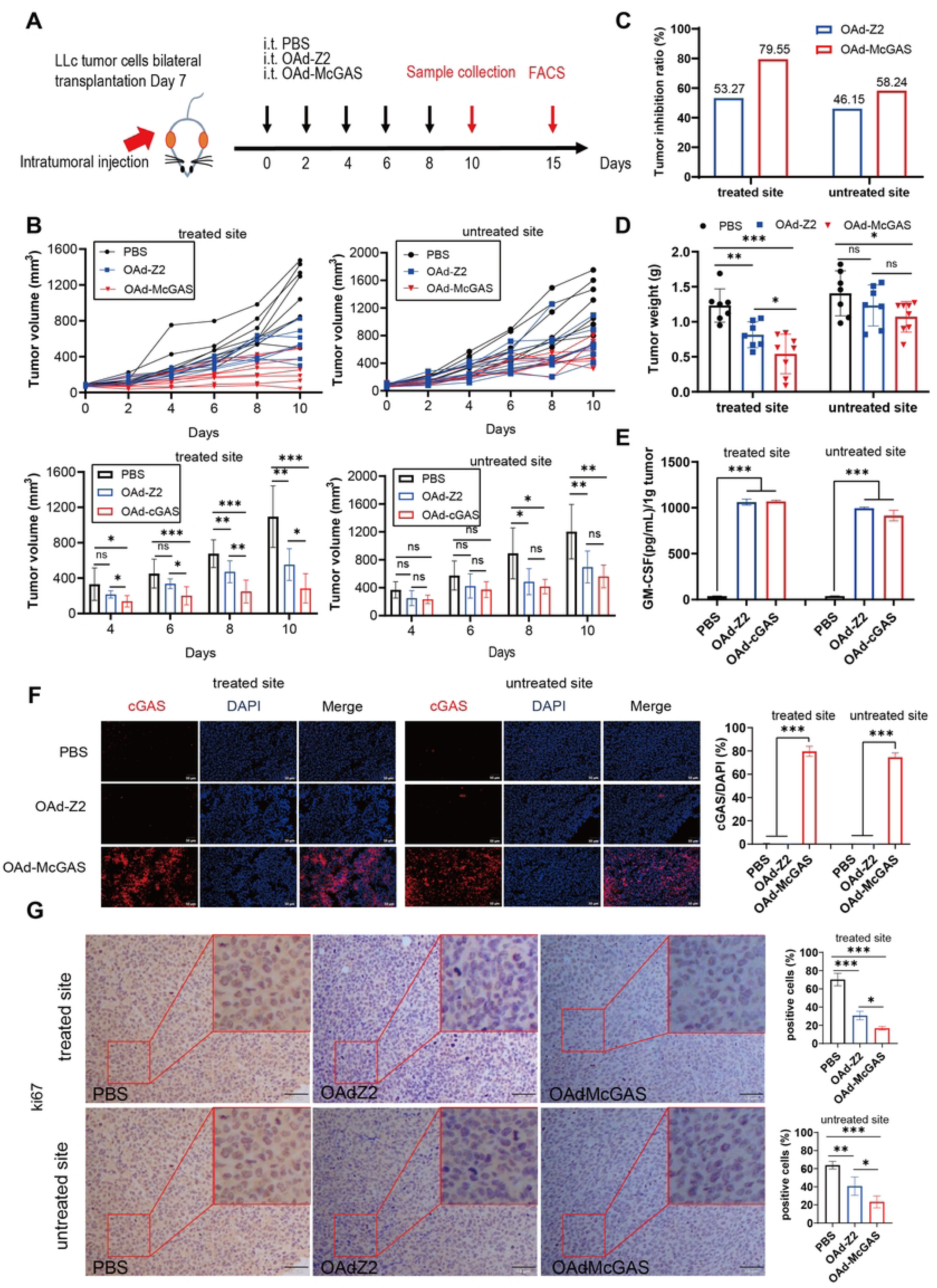
OAd-McGAS has a distant effect on tumor growth inhibition in LLC allograft tumors. **(A)** Therapeutic scheme. In brief, one side was intratumorally administered with OAds (1×10^8^ PFU/tumor) every 2 days for 5 times in the bilateral subcutaneous LLC tumor model in C57BL/6 mice, and the other side was untreated. **(B)**The growth curve of each tumor volume on the treated and untreated sides; **(C)** The tumor inhibition ratio of treated and untreated sides; **(D)** The tumor weight of treated and untreated sides; **(E)** The expression of GM-CSF in treated and untreated tumor tissues determined by ELISA; **(F)** Immunofluorescence detection of cGAS protein expression (50 μm) and quantitative analysis of cGAS^+^ area percentage in three random fields from treated and untreated tumor tissues; **(G)** Representative Ki-67 immunohistochemical staining (50 μm) and quantitative analysis of Ki-67^+^ cells in three random fields from treated and untreated tumor tissues. Data were shown as *X̄* ± *SD*, * *p* < 0.05, ** *p* < 0.01, *** *p* < 0.001, ns: no significant difference.

Although human adenoviruses exhibit variable infectivity and replication in murine cells depending on serotype and cellular context, both OAd-Z2 and OAd-McGAS effectively infected and replicated in LLC cells, as evidenced by substantial cytopathic effects and robust expression of cGAS and GM-CSF at an MOI of 200 (Fig. S2A and 1E-1F). *In vivo* analyses further revealed elevated Hexon protein levels and increased viral DNA copies in both treated and untreated tumors from OAd-treated mice compared with PBS controls (*p* < 0.05; Fig. S2B-S2C). Transgene expression of cGAS and GM-CSF was further confirmed by immunofluorescence and ELISA, respectively (*p* < 0.05; Fig. 5E-5F). Collectively, these findings indicate that OAd-McGAS exerts dual mechanisms of action in LLC tumor models: efficient infection and replication within tumor cells, coupled with functional expression of therapeutic transgenes. Hematoxylin and Eosin (H&E) staining revealed no significant tissue alterations or liver damage in mice treated with either OAd-Z2 or OAd-McGAS (Fig. S2E). Furthermore, overall body weight remained stable and gradually increased throughout the study (Fig. S2D), and i.t. administration of OAd-Z2 or OAd-McGAS did not induce severe physiological distress in the LLC transplanted tumor mouse model.

To determine whether OAd-McGAS activates the STING pathway in murine tumor tissues, tumor samples harvested on Day 10 were subjected to immunohistochemistry (IHC) staining to detect pTBK1, pIRF3, and downstream proteins, including CXCL10, IFIT1, IFIT3, TNF-α, and IFN-γ. As shown in Fig. 6A–6B and Fig. S3A–S3E, the OAd-McGAS group exhibited robust upregulation of STING signaling pathway–related proteins at both treated and untreated tumor sites (*p* < 0.05). Further supporting these findings, *in vitro* assays confirmed that OAd-McGAS enhanced the phosphorylation of STING, TBK1, and IRF3 (Fig. 6). Interestingly, higher doses of OAd-HcGAS induced weaker STING pathway activation, a paradoxical effect that may be attributed to OAd-mediated inhibition of the STING signaling pathway^[44]^.

**Figure 6.**
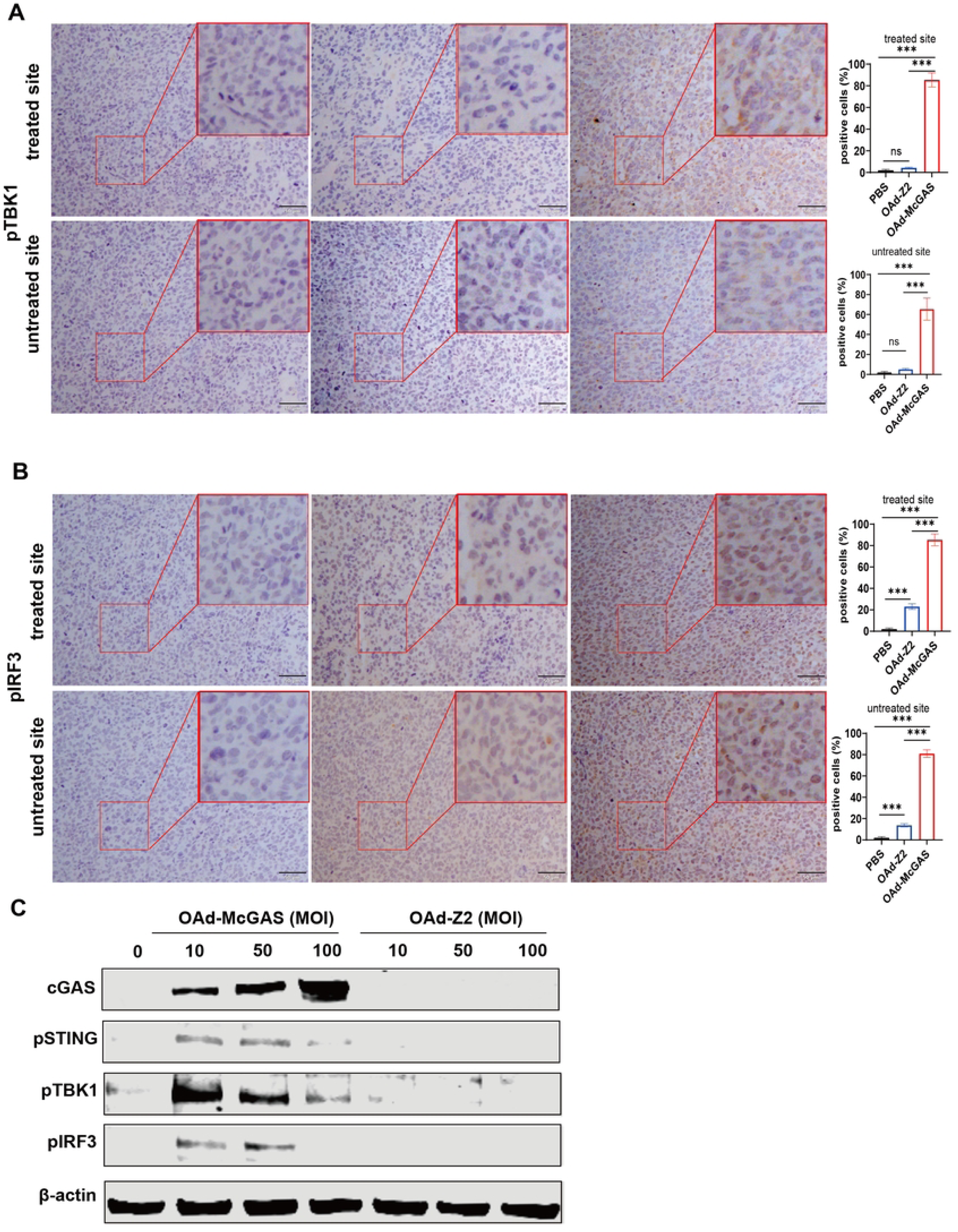
OAd-McGAS therapy can effectively activate STING pathway. Immunohistochemical staining was used to observe the status of pTBK1 (50 μm) in the tumor tissues on the treated side and the untreated side **(A)**, and the proportion of pTBK1 positive cells in the three random fields was statistically analyzed; **(B)** pIRF3 situation (50 μm) and statistics of the proportion of PIRF3-positive cells in three random fields; **(C)** Western blotting of pSTING, pTBK1, and pIRF3 in LLC cells infected with oncolytic virus at varying MOIs for 48 h. Data were shown as *X̄* ± *SD*, * *p* < 0.05, ** *p* < 0.01, *** *p* < 0.001, ns: no significant difference.

### OAd-McGAS therapy efficiently increases tumor-infiltrating lymphocytes

As a potent cytoplasmic DNA sensor, cGAS initiates STING pathway activation in tumor cells and amplifies innate immune responses. To assess OAd-mediated immune responses, tumors were harvested on Day 15 post-treatment and analyzed by flow cytometry for tumor-infiltrating lymphocytes (TILs). As illustrated in Fig. 7A-7C, Compared to the PBS and OAd-Z2 groups, the OAd-McGAS group showed an increase in the numbers of CD8⁺ T cells, conventional dendritic cells (cDCs), and tumor-associated macrophages (TAMs) within the tumor microenvironment on both the treated and untreated sides. No changes in NK cells were detected (Fig. 7D). Immunofluorescence performed on Day 10 corroborated these findings, revealing higher CD8⁺ T cell density in both treated and untreated tumors (Fig. 7E, *p* < 0.05). Further cytotoxicity assays revealed that TILs from OAd-McGAS–treated tumors effectively lysed LLC-EGFP target cells, as indicated by reduced green fluorescence and significantly lower RFU values at E:T co-culture (Fig. 7F-7G), demonstrating enhanced TIL killing activity that likely underlies its antitumor efficacy.

**Figure 7.**
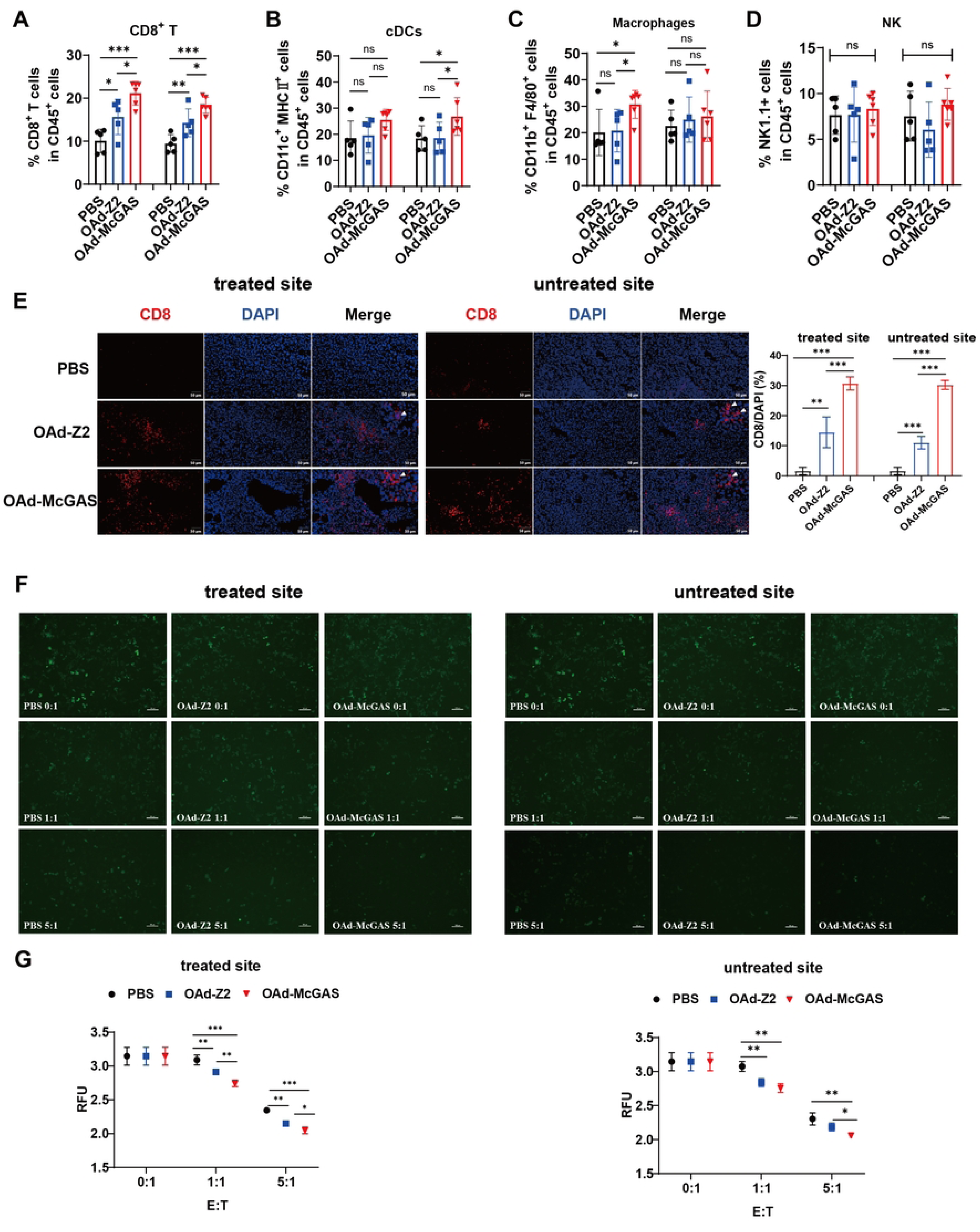
OAd-McGAS enhanced the tumoral recruitment of infiltrated lymphocytes, has obvious distance effect. LLc tumors were harvested on Day 15, digested enzymatically and stained for surface markers. FACS analysis of CD45+ CD8^+^ T cells **(A)**, CD45^+^CD3^-^CD11c^+^MHC Ⅱ ^+^ cDCs **(B)**, CD45^+^CD3^-^CD11b^+^F4/80^+^ Marophages **(C)**, CD45^+^CD3^-^NK1.1^+^ NK cells **(D)**. **(E)** Representative figures for CD8^+^ T cells, in treated and untreated tumor tissues harvested from C57BL/6 10 day after the initiation of treatment were presented by IF and quantification analysis. TILs were separated from treated or untreated tumor and cocultured with target cells at E:T ratio of 0:1, 1:1 and 5:1, and the surviving target cells were confirmed, **(F)** Treated site and untreated site, observed by fluorescence microscope. **(G)** Treated site and untreated site, recorded by microplate reader. Data were shown as *X̄* ± *SD*, * *p* < 0.05, ** *p* < 0.01, *** *p* < 0.001, ns: no significant difference.

### The combination therapy of OAd-McGAS and anti-PD-L1 demonstrates excellent therapeutic efficacy

As mentioned earlier, while OAd-McGAS is effective in treating LLc lung cancer, its therapeutic efficacy is limited and cannot completely cure the disease. Previous studies have shown that activation of the STING pathway significantly increases the expression of PD-L1 protein on tumor cells^[45]^, which our experimental data also confirm (Fig. 3H). Therefore, a combination therapy experiment was conducted as illustrated in Figure. 8A. The results demonstrated that OAd-McGAS significantly enhanced the therapeutic effect of the PD-L1 antibody. Among the five mice that received the combination treatment, three showed tumor regression (Fig. 8B-8D), with a tumor inhibition rate of 95.22% (Fig. 8E). Flow cytometry was used to prove the capacity of the combination therapy to recruit immune cells. Combination therapy with OAd-McGAS significantly increased the proportions of CD4⁺ T cells and CD8⁺ T cells in the spleens of mice treated with PD-L1 antibody (Fig. 8F, *p* < 0.05), while the proportion of NK cells remained unchanged (Fig. 8G). The results demonstrated that OAd-McGAS significantly enhances the anti-tumor efficacy of the PD-L1 antibody.

**Figure 8.**
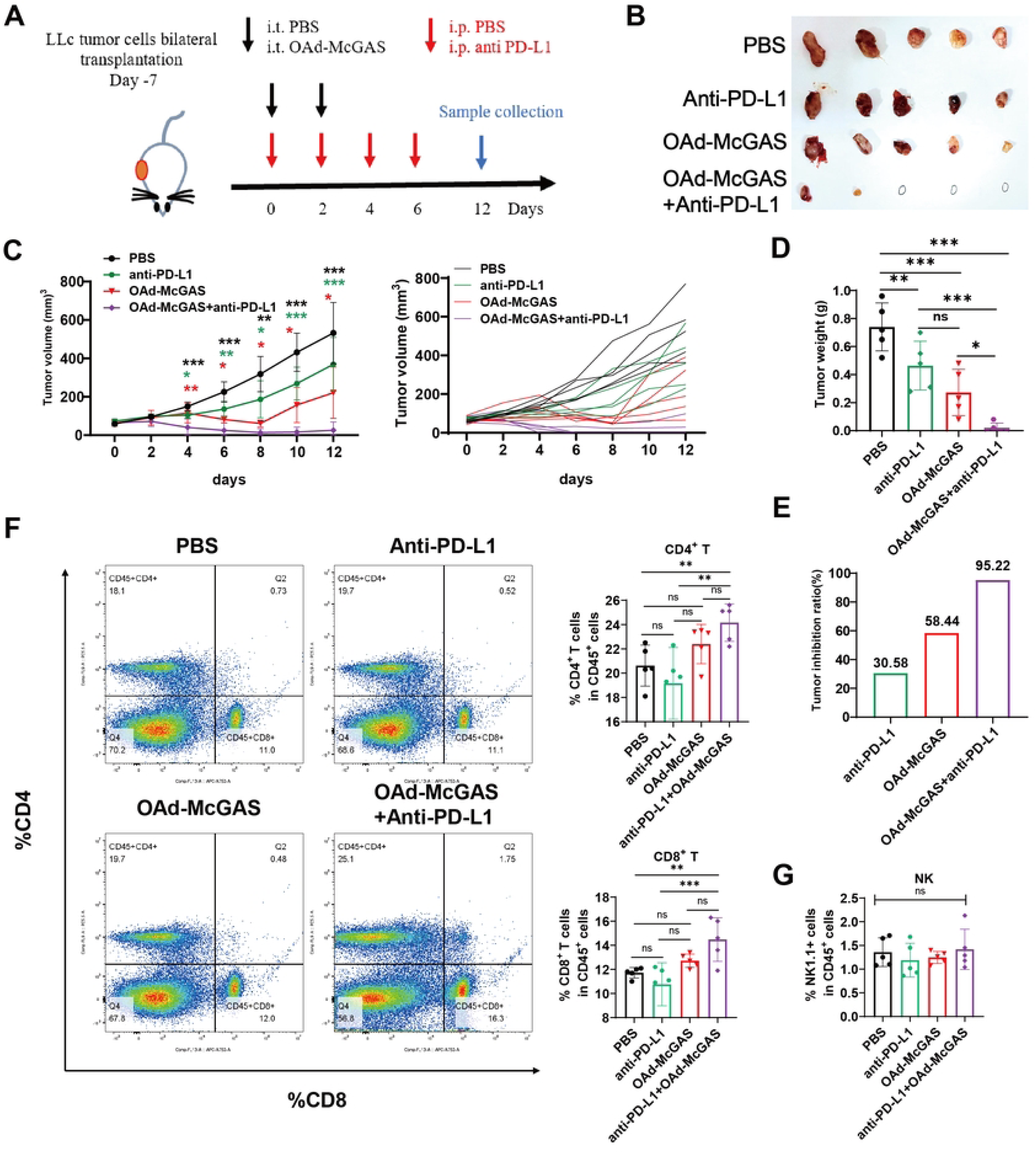
OAd-McGAS strongly improves the efficiency of anti-PD-L1 therapy. **(A)** Schematic diagram of the experimental setup for the combination therapy of recombinant adenovirus and PD-L1 antibody in LLc allograft model. OAd-McGAS were administered intratumorally (1×10⁸ PFU/tumor) every 2 days for a total of 2 doses, while PD-L1 antibody was administered intraperitoneally at a dose of 5 mg/kg every 2 days for a total of 4 doses. **(B)** Representative tumor images. **(C)** Tumor growth curves. **(D)** Tumor Weight. **(E)** Tumor inhibition ratio. FACS analysis of CD45^+^ CD4/8^+^ T cells **(F)** and CD45^+^ NK cells **(G)**. Data were shown as *X̄* ± *SD*, * *p* < 0.05, ** *p* < 0.01, *** *p* < 0.001, ns: no significant difference.

## Discussion

The cGAS-STING signaling pathway is activated when cGAS senses cytoplasmic dsDNA in tumor cells, triggering tumor-specific CD8^+^ T cell activation and adaptive immune responses, accompanied by the release of proinflammatory cytokines and chemokines^[30]^. This cascade promotes lymphocyte infiltration and modulates the immunosuppressive tumor microenvironment, positioning STING as a promising cancer therapy target. Despite its therapeutic potential, clinical application of STING agonists is hindered by limited efficacy, systemic inflammation, autoimmune-like side effects, and delivery difficulties^[32]^. Given cGAS’s pivotal role in STING activation, we hypothesized that i.t. delivery of cGAS-armed OAds could locally activate STING signaling and exert anti-tumor effects. Building on our previous hTERT-promoter–regulated Ad5/3 chimeric OAd-Z2 expressing GM-CSF, we engineered OAd-HcGAS and OAd-McGAS by P2A-linking human or murine cGAS to the E1B55k locus of the OAd-Z2 genome. This strategy generated hTERT-regulated Ad5/3 chimeric oncolytic adenoviruses, OAd-HcGAS and OAd-McGAS, co-expressing GM-CSF and cGAS. *In vitro* experiments demonstrated that OAd-HcGAS not only effectively curbed neoplastic proliferation but also potently activated the STING signaling pathway. Furthermore, in a bilateral lung cancer xenograft model, OAds carrying the cGAS gene exhibited anti-tumor effects through STING pathway engagement, enhanced lymphocyte infiltration, and synergistic suppression of tumor growth on both treated and untreated sides, which collectively validates a pronounced “abscopal effect”.

*In vitro* experiments, we validated that OAd-HcGAS activated the STING pathway in NCI-H226 cells. A key concern was whether rapid vector clearance post-STING activation would impair viral production; however, comparative replication assays in HEK 293 and NCI-H226 cells showed comparable replication capacities among H14, OAd-Z2, OAd-HcGAS, and OAd-McGAS. In conclusion, OAds encoding cGAS effectively activate the STING signaling pathway without compromising viral replication efficiency. A study by Eric Lam et al. revealed that the cGAS/STING/TBK1/IRF3 signaling cascade is not directly targeted by viral anti-host strategies, and adenovirus-induced activation of the cGAS/STING DNA response does not significantly affect viral replication efficiency^[46]^. Several mechanisms may explain this phenomenon. The adenoviral E1A protein can circumvent host antiviral responses by interacting with cellular proteins to inhibit viral DNA recognition, thereby suppressing the activities of cGAS and STING^[44, 47]^. Moreover, as adenovirus replication occurs within the nucleus^[48–51]^, cGAS primarily senses and binds exogenous or endogenous cytoplasmic dsDNA^[52]^. To avoid detection by cGAS, adenoviruses encapsulate their DNA within protein coats. During OAd construction, the E1A CR2 region, which binds to the RB protein, was deleted. Consequently, E1A is not inhibited by interferon, allowing viral replication to proceed unaffected even when OAds activate the IFN-I system. Furthermore, numerous advanced malignant tumor cells harbor intrinsic defects in the STING pathway, thereby creating an ideal environment for OVs replication^[33]^. It should be noted that the complete molecular mechanisms underlying this reverse regulation phenomenon have not yet been fully elucidated. Further exploration of STING downstream effectors related to viral replication can be conducted through proteomics profiling.

Low CAR expression in mouse LLC cells is known to compromise the infectivity of HAdV5 and other adenoviruses that primarily utilize CAR as their receptor^[53, 54]^. In this study, Ad5/3 fiber chimeric OAds were employed, though their infectivity toward LLC cells remained unclear. Consequently, LLC cells were infected with OAd-McGAS, and the results demonstrated efficient secretion of GM-CSF and expression of cGAS, along with observable CPE. Furthermore, the expression of GM-CSF and cGAS was detectable in tumor tissues from LLC xenograft mice subsequent to i.t. injection of OAd-McGAS. These results indicate that OAd-McGAS can infect LLC cells and drive the expression of target proteins.

The TME exerts a pivotal regulatory role in the therapeutic efficacy of OAds. In this study, we demonstrated that OAd-HcGAS effectively activates the STING pathway in tumor cells under *in vitro* conditions, which in turn elicits IFN-I induction, upregulates the expression of chemokines CXCL10 and CXCL9, and enhances PD-L1 expression. Using the Transwell assay, we verified that chemokines secreted by OAd-HcGAS–infected NCI-H226 cells potently recruit lymphocytes to migrate into Transwell chambers. Notably, significant activation of human T cells and NK cells was observed in an *in vitro* co-culture system established using OAd-infected tumor cells. In C57BL/6N mice bearing LLC xenografts, substantial infiltration of lymphocytes—including CD8^+^ T lymphocytes, cDCs, and tumor-associated macrophages—was detected within the tumor tissues. Furthermore, CD8+ T cells isolated from these tumor tissues exhibited remarkable cytotoxicity against LLC-EGFP target cells. Collectively, cGAS-armed OAds can reprogram the TME, transforming “cold tumors” into “hot tumors.” Leveraging this unique property—for instance, through combination therapy of OAd-HcGAS with PD-1/PD-L1 blockade—may overcome immune evasion via synergistic anti-tumor effects, thereby optimizing anti-tumor efficacy.

In both *in vitro* and *in vivo* experiments, cGAS-armed OAds demonstrated more potent anti-tumor activity than OAd-Z2, with strengthened immune-stimulatory potential and the ability to induce immunogenic cell death in tumor cells. For one, cGAS activates the STING signaling pathway, which not only stimulates innate immunity (e.g., DCs, NK cells, macrophages), but also initiates adaptive immunity (e.g., CD8^+^ T cells). This cascade reshapes the TME by critically regulating the activation and recruitment of dendritic cells, T cells, and NK cells through IFN-I signaling, ultimately mediating anti-tumor immunity^[55–59]^. For another, this enhanced activity may stem from synergistic interaction between GM-CSF and the STING pathway. However, in the absence of a control vector expressing cGAS alone, the study cannot definitively determine whether GM-CSF augments STING-mediated antitumor activity or whether the two pathways operate independently.

In C57BL/6N mice, OAds elicited a clear abscopal response in bilateral lung cancer xenografts. In contrast, no such effect was detected in BALB/c-nu mice bearing bilateral NCI-H226 tumors. These findings suggest that the abscopal activity of OAds is primarily contingent upon the activation of systemic anti-tumor immune responses^[60, 61]^. In this study, significant lymphocyte infiltration was observed in the TME, including CD8^+^ T lymphocytes, cDC cells, and tumor-associated macrophages, on both the treated and untreated sides. The robust cytotoxic capacity exhibited by these infiltrating lymphocytes further supports effective induction of systemic antitumor immunity.

We engineered oncolytic adenoviruses, OAd-HcGAS and OAd-McGAS, co-expressing cGAS and GM-CSF, which in combination with PD-L1 antibody demonstrated potent antitumor efficacy, offering a novel therapeutic strategy for lung cancer treatment. In subsequent studies, we further optimized these vectors to circumvent the constraint imposed by STING signaling loss and systematically clarified the molecular mechanisms that enable efficient viral replication following STING pathway activation. Particular emphasis was placed on the interactions between virus-encoded immunomodulatory proteins and host antiviral pathways. By delineating the distinct regulatory networks that differentiate OVs from conventional antiviral responses, these findings lay a critical theoretical foundation for the development of next-generation OV-based therapies.

## Materials and Methods

### Cell culture

The human squamous lung carcinoma cell line NCI-H226, lung adenocarcinoma cell line A549, colon cancer cell line HCT116, melanoma cell line A2058, prostate cancer cell line PC3, as well as murine Lewis lung carcinoma cell lines LLC and LLC-EGFP were obtained from the Institute of Basic Medical Sciences (IBMS) of the Chinese Academy of Medical Sciences. NCI-H226 cells were cultured in RPMI 1640 medium (EallBio, 03.4007C, China); HCT116 cells in McCoy’s 5A medium (Procell system, PM150710, China); and PC3 cells in Ham’s F-12K medium (Gibco, 21127022, USA). LLC, LLC-EGFP, A549, and A2058 cells were maintained in Dulbecco’s Modified Eagle Medium (HyClone Laboratories, SH3024301, USA). All media were supplemented with 10% fetal bovine serum (Gibco, 15630-080, Australia), 100 IU/mL penicillin, 100 μg/mL streptomycin (EallBio, F240311, China), and 2 mM L-glutamine (Thermo Fisher Scientific, 21051024, USA). All cell lines were maintained in a humidified incubator at 37℃ with 5% CO_2_.

### Oncolytic adenovirus preparation

Building on our previously developed hTERT-regulated Ad5/3 chimeric oncolytic adenovirus OAd-Z2, which expresses GM-CSF in a tumor-specific manner, we fusedeither the human or murine cGAS gene to the E1B55k locus of the OAd-Z2 genome via P2A to generate the hTERT-regulated Ad5/3 chimeric oncolytic adenoviruses OAd-HcGAS and OAd-McGAS, both capable of co-expressing GM-CSF and cGAS. Both OAd-HcGAS and OAd-McGAS were rescued and propagated in HEK 293 cells. Viral titers were measured by Adeno-X Rapid Titer kit (Clontech, Mountain View, USA) after purification using cesium chloride density gradient centrifugation, as previously described.

### Transmission electron microscopy (TEM)

The purified viruses were adsorbed onto a 200-mesh copper grid for 1 min, followed by staining with phosphotungstic acid for 1 min. The grid was subsequently placed in a storage box and allowed to dry completely before examination. Imaging was performed using a JEM-1400 electron microscope, and images were acquired using an 830.10U3 CCD camera system.

### Cytotoxicity assays

The cytotoxic effects of OAd-Z2 and OAd-HcGAS were evaluated using crystal violet and CCK-8 assays across a panel of human cancer cell lines, including NCI-H226, A549, HCT116, A2058, and PC3.

For the crystal violet assay, cells were seeded in 24-well plates at a density of 2×10⁵ cells per well and infected with viruses at MOIs of 0, 0.1, 0.5, 1, 5, and 10. At 96 h post-infection, the cells were fixed with 4% paraformaldehyde for 30 min and stained with a 0.1% crystal violet solution for 10 min. After staining, the wells were gently washed three times with PBS, and images were captured.

For the CCK-8 assay, cells were plated in 96-well plates at 1.5×10⁴ cells per well and infected at MOIs of 0, 0.01, 0.1, 1, 10, and 100. At 24, 48, 72, 96, and 120 h post-infection, 10 µL of CCK-8 solution was added to each well. The absorbance at 450 nm was measured using a microplate reader within 2 h. Cell viability was calculated relative to the mock-infected control cells (set at 100%) using the following formula:

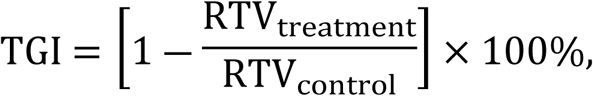

After 48 h following infection with OAd-HcGAS, NCI-H226 cells were harvested and lysed for Western blotting analysis. For PD-L1 detection, membrane proteins were extracted using a membrane protein extraction kit (Epizyme, PC202, China). Proteins were resolved under reducing conditions (100 mM 2-mercaptoethanol, boiled) on 10% SDS-PAGE gels and transferred to PVDF membranes. The membranes were then incubated overnight at 4°C with primary antibodies (detailed in the online supplemental Materials), followed by a 1 h incubation at room temperature with fluorescently labeled secondary antibody (1:10000) in the dark. Protein bands were visualized and analyzed using a two-color infrared fluorescence imaging system (ODYSSEY CLX, Li-Cor Biosciences, NE, USA). For analysis of signaling pathway proteins, the membranes were stripped and reprobed with specific antibodies. *Assessment of cytokine expression by ELISA*

For cell culture supernatants, samples were centrifuged at 2,000 × g for 5 min to remove cellular debris. The clarified supernatants were then analyzed using respective ELISA kits according to the manufacturers’ protocols for the detection of human GM-CSF, mouse GM-CSF, human IFN-β, CXCL10, CXCL9, and TNF-α.

For tumor tissue samples, approximately 0.2 g of tissue was homogenized in RIPA lysis buffer, followed by centrifugation at 12,000 rpm for 10 min. The resulting lysates were subsequently assayed for mouse GM-CSF levels using a specific ELISA kit.

### Co-culturing of immune cells and NCI-H226 cells in the presence of OAd-Z2 or OAd-HcGAS

For direct *in vitro* co-culture experiments, NCI-H226 cells were plated with PBMCs at a 1:10 ratio in RPMI-1640 medium supplemented with 10% FBS. After 24 h of incubation, the cultures were infected with OAds or OAd-HcGAS at a multiplicity of infection of 10 and incubated for an additional 48 h. The supernatant was collected for LDH detection, and the isolated PBMCs were detected by flow cytometry.

### Detection of PBMC cytotoxicity

The cytotoxicity of PBMCs was estimated by measuring LDH activity in the culture medium using the LDH Cytotoxicity Assay Kit (Beyotime, C0017, China). Briefly, assays were carried out in 96-well plates with a final sample volume of 120 µL/well. The LDH released reagent was added at 10% of the volume of the original culture medium.

### Flow cytometry analysis of P-PBMCs (CD8^+^, CD56^+^, CD69^+^ and IFN-γ^+^)

PBMCs isolated from the co-culture system were transferred into U-bottom 96-well plates and blocked with Human TruStain FcX^TM^ (BioLegend, 422302, USA). CD8-, CD56-, and CD69-conjugated fluorescent antibodies were added, and cells were incubated in the dark at 4°C for 30 min. Following a single wash, cells were fixed with fixation buffer for 20 min in the dark. Fixed cells were then resuspended in diluted Intracellular Staining Permeabilization Wash Buffer (BioLegend, 421002, USA), and centrifuged at 350 × g for 5–10 min; this step was repeated twice. Finally, IFN-γ-conjugated fluorescent antibodies were added and incubated in the dark at 4°C for 30 min.

### Animal studies

Specific pathogen-free female C57BL/6 mice, aged 6–8 weeks, were purchased from Vital River Laboratory Animal Technology Ltd. (Beijing, China) and kept under SPF conditions. All animal experiments were approved by the Animal Welfare and Research Ethics Committee of Beijing Laboratory Animal Research Center (BLARC-LAWER-202407009).

Subcutaneous tumor dimensions were measured every 2 days using a digital vernier caliper. The tumor volume (mm^3^) was calculated as L × W^2^/2, where L and W represent tumor length and width, respectively. Once tumor volumes reached approximately 50-100 mm³, mice were randomly assigned to treatment groups and received intratumoral (i.t.) injections. The tumor growth inhibition rate (TGI) was calculated based on relative changes in tumor volume using the following formula:

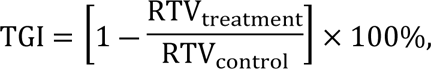

where relative tumor volume (RTV) is defined as V_t_/V_0_, with V_t_ representing the mean tumor volume at endpoint (Day 10) and V_0_ representing the mean tumor volume at baseline (Day 0).

### Multicolor flow cytometry

Tumor tissues or spleen were collected from mice, minced, and digested in RPMI-1640 medium containing 0.1 mg/mL collagenase I (Gibco, 17018029, USA), 1 mg/mL collagenase IV (Gibco, 17104019, USA) and 200 U/mL DNase I at 37°C for 1 h. The resulting cell suspension was passed through a 70-μm cell filter (Beyotime, FSTR070, China), washed with PBS, and resuspended in cell staining buffer (BioLegend, 420201, USA). Single-cell suspensions were first blocked with TruStain FcX™ (BioLegend, 101320, USA) for 10 minutes at 4°C, followed by staining with a fluorescent antibody cocktail (Supplemental Materials) for 40 minutes at 4°C. Prior to surface staining, cell viability was assessed using the Zombie NIR™ Fixable Viability Kit (BioLegend, 423106, USA) by incubating cells at room temperature in the dark for 15 minutes.

Samples were acquired on a FACSCanto II system or FACSFortessa system (BD Biosciences, CA, USA), and data were analyzed using FlowJo software (BD Biosciences, CA, USA).

### Killing capacity of TILs

CD4^+^ and CD8^+^ TILs were isolated from both treated and untreated tumor tissues using CD4/CD8 (TIL) MicroBeads (Miltenyi, 130-116-480, German). TILs were co-cultured with LLC-EGFP target cells at different effector-to-target ratios (0:1, 1:1, and 5:1) in a 37°C, 5% CO_2_ incubator for 24 h. Surviving target cells expressing green fluorescence were visualized and captured using a fluorescence microscope. Relative fluorescence units (RFU) were measured at an excitation wavelength of 479 nm and an emission wavelength of 517 nm using a microplate reader.

### Quantitative PCR (qPCR)

Total DNA was extracted from tumor tissues using a Blood/Cell/Tissue DNA Extraction Kit (Tiangen, DP304-03, China). qPCR was performed using a fluorescence-quantitative PCR kit (Tiangen, FP205-02, China) on a q225 system (KUBO technology, Beijing, China), according to the manufacturer’s instructions. The primer sequences targeting the viral hexon gene were as follows: forward, 5′-GGTGGCCATTACCTTTGACTCTTC-3′; reverse, 5′-CCACCTGTTGGTAGTCCTTGTATTTAGTATCATC-3′.

### Hematoxylin and Eosin (H&E) staining, immunohistochemistry (IHC), and immunofluorescence (IF)

Liver tissues were processed for H&E staining following standard protocols and analyzed under a light microscope (Olympus, Tokyo, Japan). For all staining procedures, tissue sections were deparaffinized, rehydrated, and washed in 1% PBST. For IHC, the sections were soaked in 3% hydrogen peroxide to block endogenous peroxidase activity and incubated with primary antibodies overnight at 4°C. The sections were then incubated with HRP-linked anti-rabbit antibodies for 20 min, developed with diaminobenzidine substrate, and counterstained with hematoxylin. For IF, tissue sections were incubated with primary antibodies overnight at 4°C, followed by fluorescently labeled secondary antibodies for 40 min the next day, and then counterstained with DAPI for 8 min. Images were acquired by fluorescence microscopy (Olympus, Tokyo, Japan) and analyzed using ImageJ software. A complete list of primary antibodies is provided in the supplemental Materials. The secondary antibodies used were as follows: for IHC, a rabbit two-step detection kit (ZSGB-Bio, PV-6001, China); for IF, Anti-Mouse Cy3 (Beyotime, P0193, China) and Anti-Rabbit FITC (Beyotime, P0186, China).

### Statistical analysis

Statistical analysis was performed using the SPSS software version 21 (Chicago, IL, USA), and one-way analysis of variance with SNK-q post-test was used to compare the means of multiple groups. *P* < 0.05 was considered significant. The data were presented as mean ± SD.

## Ethics Statement

The animal study was approved by Beijing Laboratory Animal Research Center (BLARC-LAWER-202407009). The study was conducted in accordance with the local legislation and institutional requirements.

## Consent for publication

Not applicable

## Data Availability declaration

Data sharing is not applicable to this article as no datasets were generated or analysed during the current study.

## Acknowledgments

This work was supported by National Key Research and Development Program of China (2023YFC2307900), the National Natural Science Foundation of China (32370994) and Beijing Natural Science Foundation (L222074).

## Author contributions

All authors contributed to the experimental work. Specifically, Wang Qingwen, Fu Yuanhui, and Zheng Yanpeng were responsible for the experimental design, data analysis, figure preparation, and manuscript writing. All authors have read and approved the final manuscript.

